# Mycobacterial lipoarabinomannan negatively interferes with macrophage responses to *Aspergillus fumigatus in-vitro*

**DOI:** 10.1101/2024.11.18.623945

**Authors:** LE Gonzales-Huerta, TJ Williams, R Aljohani, B Robertson, CA Evans, DPH Armstrong-James

**Affiliations:** Department of Infectious Diseases, Imperial College London, SW7 2BX, UK; Carrera de Medicina Humana, Facultad de Ciencias de la Salud, Universidad San Ignacio de Loyola, Lima 15024 Peru; Departamento de Investigación, Instituto de Medicina Traslacional, Lima 15072, Peru; Innovación Por la Salud Y Desarrollo (IPSYD), Asociación Benéfica PRISMA, Lima, 15073, Peru; IFHAD: Innovation For Health And Development, Laboratorio de Investigación y Desarrollo, Universidad Peruana Cayetano Heredia, Lima 150135 Peru; IFHAD: Innovation For Health And Development, Department of infectious disease, Imperial College London, London, UK

**Keywords:** Aspergillus fumigatus, Pulmonary Aspergillosis, tuberculosis, Mycobacterium tuberculosis, Mycobacterium smegmatis, lipoarabinomannan

## Abstract

**Introduction:** Over 1 million people have chronic pulmonary aspergillosis (CPA) secondary to pulmonary tuberculosis. Additionally, *Aspergillus fumigatus* (*Af*) has been reported as one of the most common pathogens associated with mycobacteria in patients with cystic fibrosis. Mycobacterial virulence factors, like lipoarabinomannan, have been shown to interfere with host’s intracellular pathways required for an effective immune response, however, the immunological basis for mycobacterial-fungal coinfection is still unknown. We therefore investigated the effect of lipoarabinomannan on macrophage responses against *Af*.

**Methods:** Bone marrow-derived macrophages (BMDMs) were stimulated with non-mannose-capped lipoarabinomannan (LAM) from *Mycobacterium smegmatis* or mannose-capped lipoarabinomannan (ManLAM) from *Mycobacterium tuberculosis* for 2 hours and then infected with swollen *Af* conidia. Cell death was assessed by lactate dehydrogenase release. Cytokine release was measured in supernatant using Enzyme Linked Immuno-Sorbent Assay (ELISA). Colony forming units counting and time-lapse fluorescence microscopy was performed for studying conidia killing by macrophages.

**Results:** BMDMs stimulated with LAM showed increased cell death and inflammatory cytokine release in a dose-dependent manner, characterised by a significant increase of IL-1β release. Time-lapse fluorescence microscopy and CFUs revealed that both LAM and ManLAM significantly decrease the capacity of macrophages to kill *Af* conidia within the first 6 hours of infection.

**Conclusions:** The mycobacterial virulence factor, lipoarabinomannan, disrupts macrophage capacity to efficiently clear *Af* at early stages of infection *in-vitro*.

## INTRODUCTION

*Aspergillus fumigatus* (*Af*) is a ubiquitous airborne fungus, and approximately 100-1000 conidia are believed to reach the lower respiratory tract and alveoli each day. Immunocompetent hosts clear fungi in the airway with effective innate immune responses (1). However, immunocompromised individuals, such as patients with neutropenia, cancer, lung transplant recipients, or with high exposures to corticosteroids, are susceptible to develop pulmonary aspergillosis (PA) (2). Additionally, patients with conditions that cause structural damage to the airway and pulmonary parenchyma, such as cystic fibrosis (CF), sarcoidosis, and lung cavitation post-pulmonary tuberculosis, present a higher risk for PA (3), which has an estimated 5-year mortality rate between 50% and 85% (4). In patients with CF, the association between mycobacteria and *Af* indicates increased mortality (5–7). Furthermore, the burden of PA post-tuberculosis is considered to be substantial (8), however, the causative factors for a close association between these pathogens are still unknown.

The alveolar macrophages are considered the first line of defence against *Af*. In the alveoli, macrophages take over the task of recognizing and process *Af* conidia. A protective rodlet protein layer in conidia covers the pathogen-associated molecular pattern (PAMPs) from pattern recognition receptor (PRRs) in immune cells. However, lung tissue and the body temperature of the host provide the necessary conditions for swelling and germination to initiate. The conidia swelling breaks the rodlet layer and exteriorizes galactomannan, chitin, α-glucan, and β-glucan. These components will bind to a diversity of receptors in the macrophage, inducing a mechanism that will kill conidia or inhibit their germination (9). Recognition of β-glucan by TLR-2, induces the plasma membrane receptor to activate the myeloid differentiation primary response 88 (MyD88) downstream to NF-κβ (10). Similarly, TLR-4 recognizes mannan, contributing to MyD88-mediated signalling pathway. Both, lead to increased expression and release of inflammatory cytokines, TNF-α and IL-12 (11). Additionally, C-type lectin receptors (CTL) recognize carbohydrate ligands, intervening in phagocytosis, and collaborating in the release of inflammatory cytokines (12,13). Additionally, Dectin-1 is crucial in the phagocytosis process of conidia, by recognizing β-glucans in *Af*, it activates the immunoreceptor tyrosine-based activation motif (ITAM). This leads to subsequent triggering of Src/Syk-dependent signalling pathway to contribute in the activation of NF-κβ and NFAT (14,15). Mannose receptor recognizes chitin, internalize it, and has been shown to modulate macrophage activation by increasing the production of IL-10, which dampens the inflammation (16).

NLRs have been also associated with the immune response against *Af*. Although, NOD2 expression contributes to the inflammatory response through activation of NF-κβ in *Af* infection(17), the genetic deficiency of NOD2 has been shown to be protective against invasive aspergillosis (18). NLRP3 has been reported to respond effectively to hyphal fragments by triggering the inflammasome assembly, with subsequent caspase-1 activation and IL-1β release, which requires K^+^ efflux and ROS production (19). Muramyl dipeptide (MDP) is a well-known bacterial ligand for NOD2 (20), but it is absent in *Af*. It has been suggested that inflammasome assembly is triggered by nucleic acid structures released by *Af* (21,22).

Lipoarabinomannan is one of the most important virulence factors in mycobacteria. It is constituted by a polymannosylated phosphatidylinositol core with an arabinan moiety. This arabinan termini can be capped with an inositol phosphate residue or with mannose, depending on the mycobacterial specie, and it modifies its immunogenicity (23). Mannose-capped lipoarabinomannan (ManLAM) produced by *Mycobacterium tuberculosis*, has been shown to interfere with normal phagolysosome function, inflammatory cytokine production, and to inhibit apoptosis (24–26). Alternatively, the lipoarabinomannan produced by *Mycobacterium smegmatis*, lacks the mannose cap and has been showed to induce a strong TLR-2 mediated immune response, while it decreases phagocytosis (27).

Epidemiological and clinical data suggest that mycobacteria infection potentially facilitates fungal colonization and respiratory disease. Moreover, mycobacterial lipoarabinomannan has been shown to interfere with the cellular mechanisms that are required for an effective response to *Af*. Therefore, we hypothesize that mycobacterial lipoarabinomannan negatively modulates macrophage responses to *Af*.

## MATERIAL AND METHODS

### Culture and preparation of *Aspergillus fumigatus* (*Af*)

*Af* strains CEA-10 (FGSC A1163) was obtained from the Fungal Genetics Stock Center and the American Type Culture Collection (ATCC). A GFP-expressing *Af* derived from ATCC46645 strain (ATCC46645-eGFP) was kindly given by Frank Ebel. A dsRED-expressing *Af* (AF293) was a kind gift from Georgios Chamilos. *Af* was cultured in Sabouraud dextrose agar (Oxiod® Cat. CM0041B) for 3-4 days at 37°C and harvested by washing with DPBS + 0.1% Tween-20. Conidia suspension was filtered using sterile Miracloth™ (Millipore™, Cat. 475855). For swollen conidia, 10^6^ conidia per mL of RPMI-1640 (Gibco® Cat. 11875-093) were incubated in petri-dish at 37°C + 5% CO_2_ for 3 hours.

### Tracking of conidia swelling and germination

Resting *Af* conidia was harvested and resuspended at a concentration of 5 × 10^5^ in RPMI-1640. LAM (BEI resources, Cat. NR-14849) and ManLAM (BEI resources NR-14848) were added to the media at a concentration of 1 μg/mL. *Af* conidia were transferred to a 96-well plate (Greiner® Cat. 655090) at a concentration of 10^5^ per well in 200 μL.Conidia were imaged in a widefield inverted microscope (Zeiss® Cell Discoverer 7), with a heated chamber at 37°C and 5% CO_2_. Cells were imaged every 15 min for 12 hours in a single z-plane.Image analysis was performed using FIJI. Nine conidia, properly focused for at least 9 hours, were selected from each FOV. Manual measurements of conidial diameter were taken for every hour of incubation until point of germination was reached.

### Bone marrow-derived macrophages (BMDMs)

Healthy C57BL/6J mice between 8 and 12 weeks old were euthanized with cervical dislocation. Secondary confirmation was performed with dissection of the femoral artery. Bone marrow was flushed out of mice femurs with cold PBS and filtered through a 40 μm cell strainer. Cells were centrifuged and resuspended in 60 mL of cRPMI + 40 ng/mL m-CSF (PeproTech® Cat. 315-02) + 50 μg/mL Gentamicin (Gibco®, Cat. 11540506). Cells were cultured in petri-dishes at 37°C + 5% CO_2_. After 3 days, additional 5 mL of fresh media was added, and cells incubated for 4 more days to complete differentiation. To recover BMDMs, medium was discarded, replaced with cold PBS + 2 mM EDTA (Millipore® Cat. 324506-100ML) and petri-dishes placed in the fridge for 20min. For experimentation, macrophages were seeded in a 96-well flat bottom plate at a concentration of 5 × 10^4^ per well in 200 μL of cRPMI + 20 ng/mL m-CSF. Cells were activated with overnight with 200 U/mL of IFN-γ (Peprotech® Cat. 315-05). On the day of infection, IFN-γ was removed and cells were stimulated with lipoarabinomannan (BEI resources, Cat. NR-14849 and NR-14848) for 2 hours. Cells were infected with CEA-10 *Af* or dsRED-*Af* swollen conidia at MOI=2.

### LDH assay

LDH release was tested using the Cytotox-96® Non-Radioactive Cytotoxicity Assay (Promega® Cat. G1780) according to manufacturer’s instructions. 50 μL of fresh supernatant was employed.

### Cytokine release

BMDMs, were seeded at 5 × 10^4^ cells per well and treated for 2 hours with LAM or ManLAM [0.1, 0.5, and 1 μg/mL], prior to infection with CEA-10 Af swollen conidia at MOI=2. Supernatant was collected 24 hpi.

Cytokines were studied using the supernatant from tissue culture and a sandwich ELISA kit from Bio-Techne for TNF-α (R&D Systems Cat. DY410), IL-1β (R&D Systems Cat. DY401) and CXCL1/KC (R&D Systems Cat. DY453) according to manufacturer’s instructions in Nunc MaxiSorp™ 96-well plates (Invitrogen™, Cat. 44-2404-21).

### Colony forming units (CFUs)

10^5^ cells per well were prepared and treated in a 24 well plate for experimentation. Cells were infected with CEA-10 swollen conidia at MOI=2. At 3 hpi, supernatant was removed, cells were washed with warm PBS and lysed with 500 μL of DPBS (Gibco® Cat. 14190-136) + 0.1% Tween-20 (Sigma-Aldrich®, Cat. 9416-100ML). Serial dilutions were prepared from lysed cells. 100 μL of lysed cells dilution (1:100) was transferred and spread on a 9 mm petri-dish containing 15 mL of Sabouraud agar (Oxiod® Cat. CM0041B), incubated at 37°C for 18-24 hours, and counted manually.

### FLARE conidia

Fluorescent Aspergillus Reporter (FLARE) conidia was prepared according to the protocol published by Tobias Hohl’ group (28). dsRED-*Af* aliquots were centrifuged at 9300 g for 10 min and the supernatant discarded. The pellet is resuspended in 1 mL of 0.05 M NaHCO_3_ pH 8.3 + 500 μg of biotin-XX, SSE (Invitrogen, Cat. 11524197) and incubated at 4°C for 2 h. Next, *Af* is centrifuged, and pellet resuspended in 1 mL of 100 mM Tris-HCl pH 8.0 for 1 h at room temperature. Then, *Af* is centrifuged and Tris-HCl discarded. Pellet is resuspended in 1 mL PBS with 20 μg/mL of Streptavidin-AF633 (Invitrogen, Cat. S21375) and incubated for 40 min at room temperature. Conidia is then centrifuged at 9300 g x 10 min, supernatant removed, and pellet resuspended in PBS. For swollen conidia, fungi are counted, new aliquots of 2×10^6^ or 4×10^6^ conidia per mL are prepared with RPMI-1640 (Gibco® Cat. 11875-093) and incubated at 37°C +5% CO_2_ for 3 h. Finally, conidia are centrifuged again, medium discarded, and pellet resuspended in culture medium.

### Microscopy tracking of conidia killing assay

Macrophages were seeded in a 96-well (Greiner® Cat. 655090) at a concentration of 5 × 10^4^ per well for BMDMs. On the day of infection, IFN-γ and m-CSF was removed to stimulate the macrophages with 1 μg/mL of LAM, ManLAM, in 100 μL of cRPMI without phenol red. After 1 hour, 100 μL of medium with 100 nM of LysoTracker™ Blue DND-22 (Invitrogen™ Cat. L7525) was added to each condition, preserving the mycobacterial ligands’ concentration. After 2 hours of stimulation, the medium was removed and replaced with 200 μL of fresh cRPMI + ligands. Next, 50 μL of medium with 10^5^ FLARE swollen conidia was added. For controls, 50 μL of medium without *Af* conidia is added to get all condition to 250 μL.

For BMDMs, cells were imaged in an automated high-content widefield inverted microscope (Zeiss® Cell Discoverer 7), with a heated chamber at 37°C and 5% CO_2_. One FOV per well was set for time-lapse images, acquired with Zen Blue 3.1 software in 16-bit. Cells were imaged every 15min for 6 hours in a single z-plane.

Image analysis was performed using FIJI. Channels for Lysotracker™ Blue DND-22, dsRED and AF633 were split using a FOV from an unstimulated well with *Af* infection to manually determine the threshold for positivity. These threshold values were set for the rest of FOVs. dsRED and AF633 channels were employed for counting conidia killing manually.

### Statistical analysis

All values are shown as biological replicates. All statistical analysis were performed on biological replicates. To determine statistical significance between two unmatched groups Students-T test was used. For significance testing in data normally distributed between more than two groups in one variable, One-way analysis of variance (ANOVA) was applied. Comparisons between two or more groups and two variables were analysed using Two-way ANOVA. Statistical analysis was perform using GraphPad Prism 9.4.1.

## RESULTS

### Lipoarabinomannan does not alter conidial swelling and germination

We investigated if lipoarabinomannan alters conidial swelling and the incubation time required for germination. No statistically significant differences were found between conditions. Lipoarabinomannan did not change the swelling rate or time of incubation for germination (Figure 1 A, B). Mean diameter for unstimulated CEA-10 conidia was 4.4 µm at 0 hours and reached 7.3 µm by 6 hours. For dsRED-*Af*, the diameter ranged from 4.5 µm at 0 hours to 7.9 µm at 6 hours.

**Figure 1.**
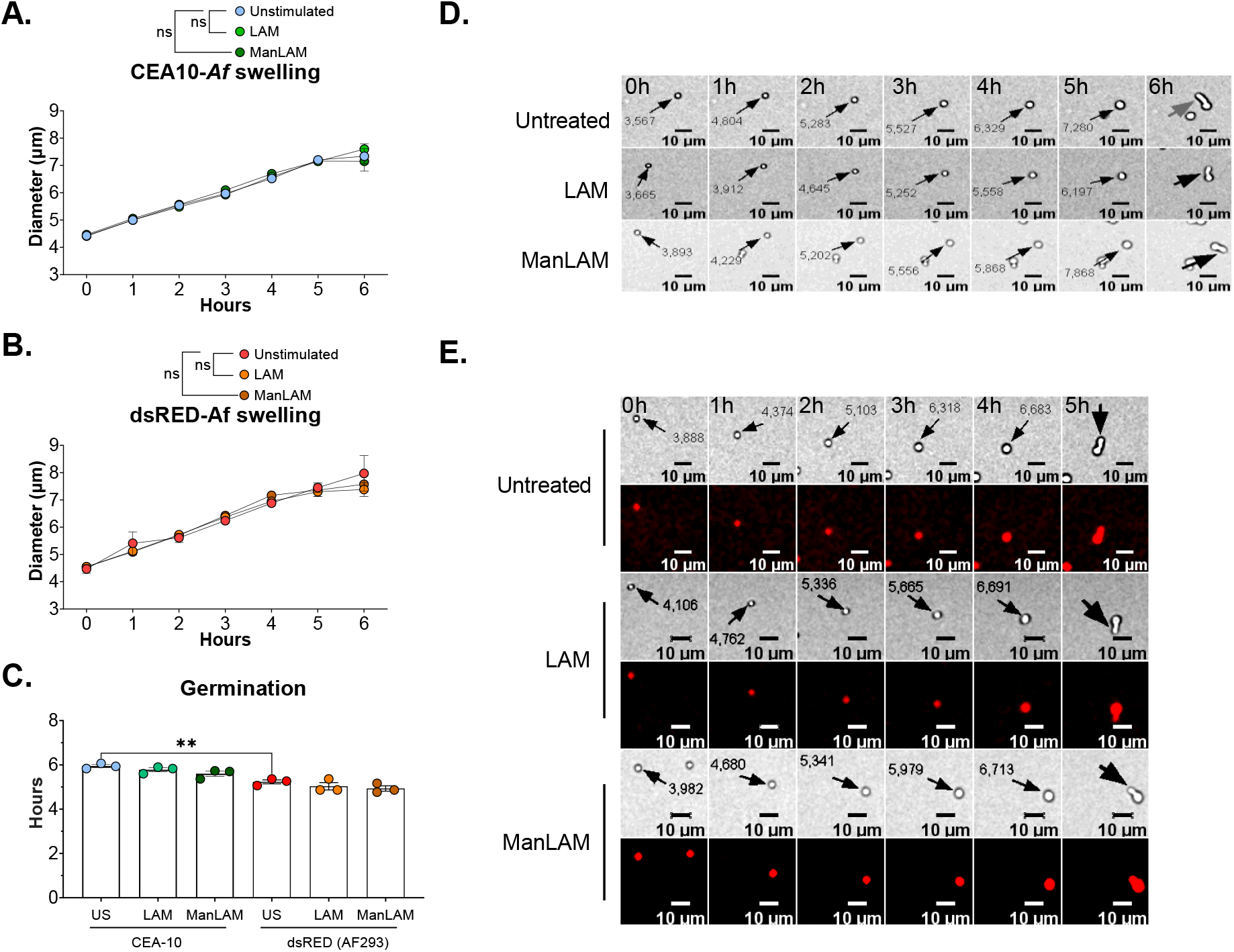
Lipoarabinomannan does not interfere with *Af* fitness. **A**. Conidial diameter of CEA-10 *Af* strain over time. **B**. Conidal diameter for ds-RED *Af*. **C**. Comparison of the time of incubation at 37°C +5% CO_2_ required to reach germination. **D**. Representative images of CEA-10 strain swelling until it reaches germination. **E**. Representative images of dsRED strain swelling and germination. Statistically significant differences were tested with two-way ANOVA for time-lapse microscopy experiment and one-way ANOVA for germination data. Statistical tests were performed on results from 3 experimental replicates (n=3). Error bars in SEM. p value * < 0.05, ** < 0.01, *** < 0.001, **** < 0.0001.

For CEA-10, germination started approximately at 5.9 hours of incubation. LAM stimulation averaged 5.8 hours, and ManLAM 5.6 hours. Similar results were obtained with dsRED-*Af*, with unstimulated conidia reaching germination stage at 5.2 hours. For LAM, germination time-point was reached at 5 hours, and for ManLAM at 4.9 hours. These were not statistically significant differences (Figure 1 C).

### LAM increases cell death in BMDMs infected with *Af*

LAM showed to increase cell death in BMDMs in a dose-dependent manner by 12 hpi. Mean values for cell death were 27%, 35.9%, 49.9%, and 54.9% for unstimulated, 0.1 µg/mL, 0.5 µg/mL, and 1 µg/mL of LAM respectively. The statistically significant difference is lost by 24 hpi; however, a clear trend persists with higher mean values for macrophages stimulated with LAM (Figure 2). ManLAM did not produce any statistically significant differences.

**Figure 2.**
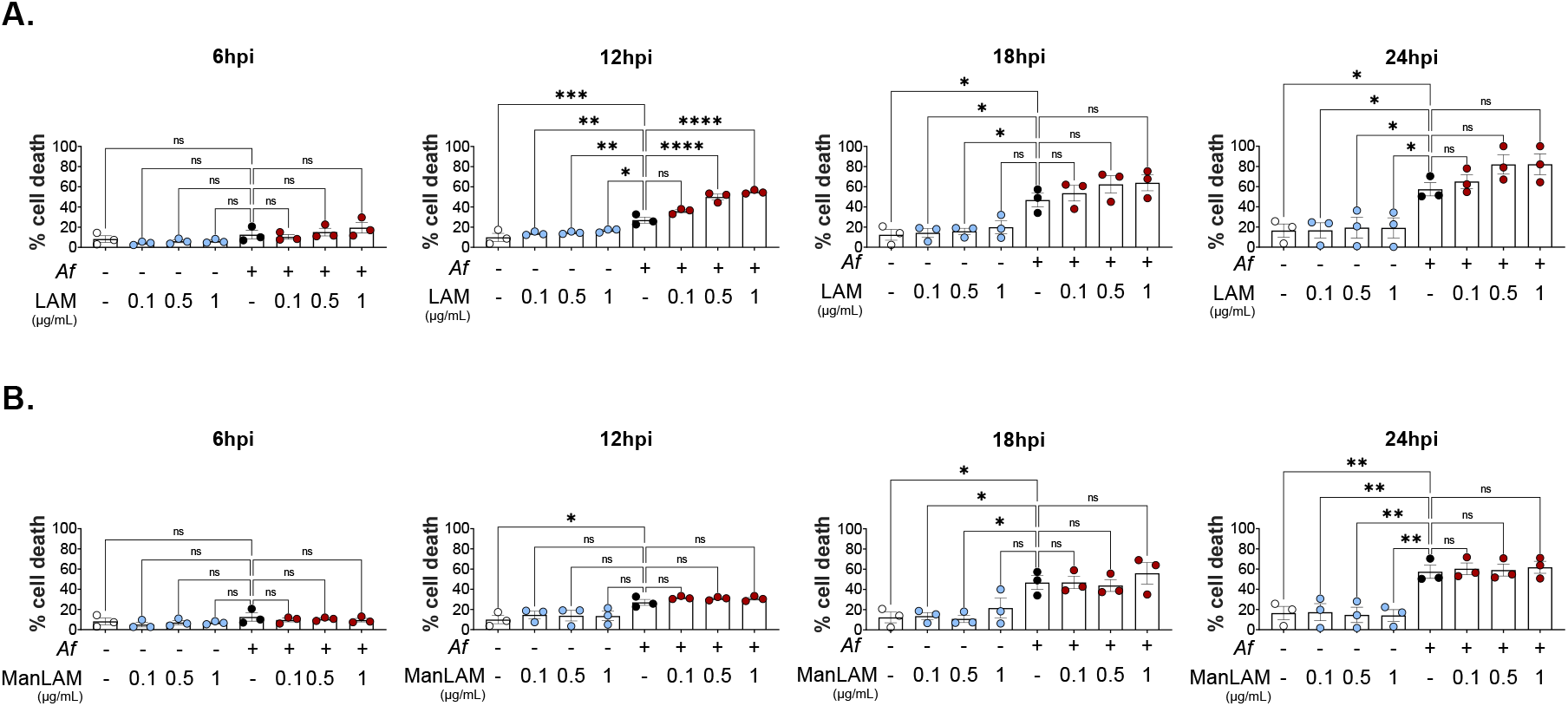
LAM increases cell death in *Af* infection. **A**. Cytotoxicity at 6, 12, 18 and 24 hpi measured as LDH release. Three concentrations of LAM (0.1µg/mL, 0.5µg/mL, and 1µg/mL) were tested in uninfected controls (blue) and infected with swollen conidia of CEA-10 *Af* (red). **B**. Cytotoxicity at 6, 12, and 24 hpi with ManLAM at 0.1 µg/mL, 0.5 µg/mL, and 1 µg/mL. One-way ANOVA was performed on biological replicates. Error bars in SEM.

### LAM increases IL-1β release from *Af* infected BMDMs

BMDMs stimulated with LAM produced a significant increase of IL-1β release. Mean value for cells stimulated with 1 µg/mL of LAM was 385.8 pg/mL, whereas for unstimulated cells was 16.71 pg/mL (Figure 3). It is important to note that at 24 hpi, more than 80% of LAM-stimulated macrophages had died, but only 57% of unstimulated macrophages were dead at 24 hpi. However, the increasing trend in IL-1β is suggestive of a stronger inflammatory response. Although there was an increasing trend in CXCL1 with higher concentrations of LAM, it was not statistically significant. TNF-α did not show a statistically significant difference. Finally, ManLAM did not have any effect on BMDMs cytokine release.

**Figure 3.**
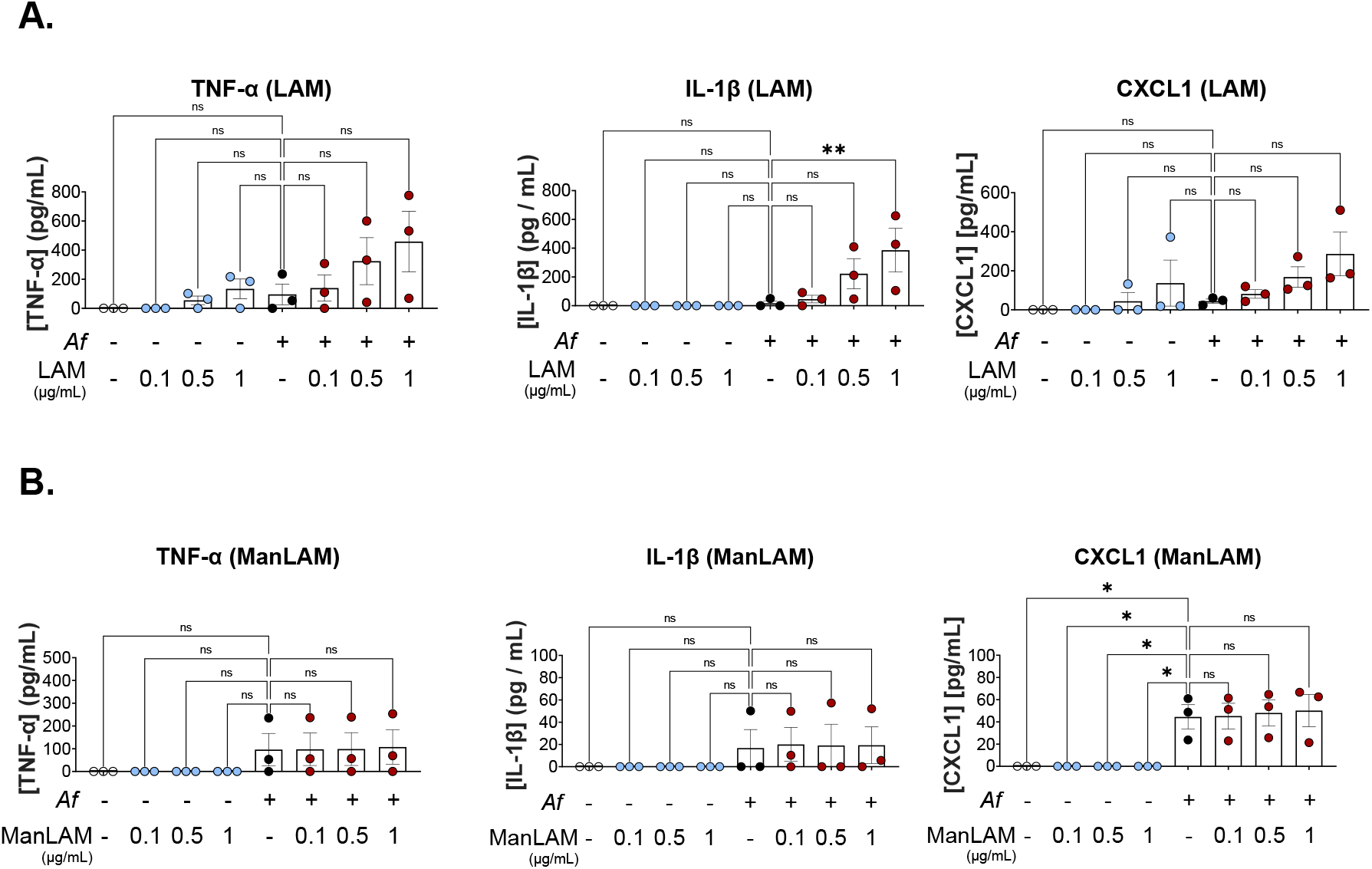
LAM increases IL-1β release with *Af* infection. **A**. Inflammatory cytokine release from BMDMs stimulated with different concentrations of LAM at 24 hpi. **B**. Inflammatory cytokine release from BMDMs stimulated with different concentrations of ManLAM at 24 hpi. One-way ANOVA was performed on biological replicates. Error bars in SEM.

### Mycobacterial lipoarabinomannan reduces BMDMs *Af* killing rate

Next, we studied the effect 1 μg/mL for LAM and ManLAM on lysosome delivery and conidia killing. Pixels colocalization between LysotrackerBlue and dsRED-expressing conidia showed no statistically significant differences between lipoarabinomannan and unstimulated cells(Figure 4 A).

**Figure 4.**
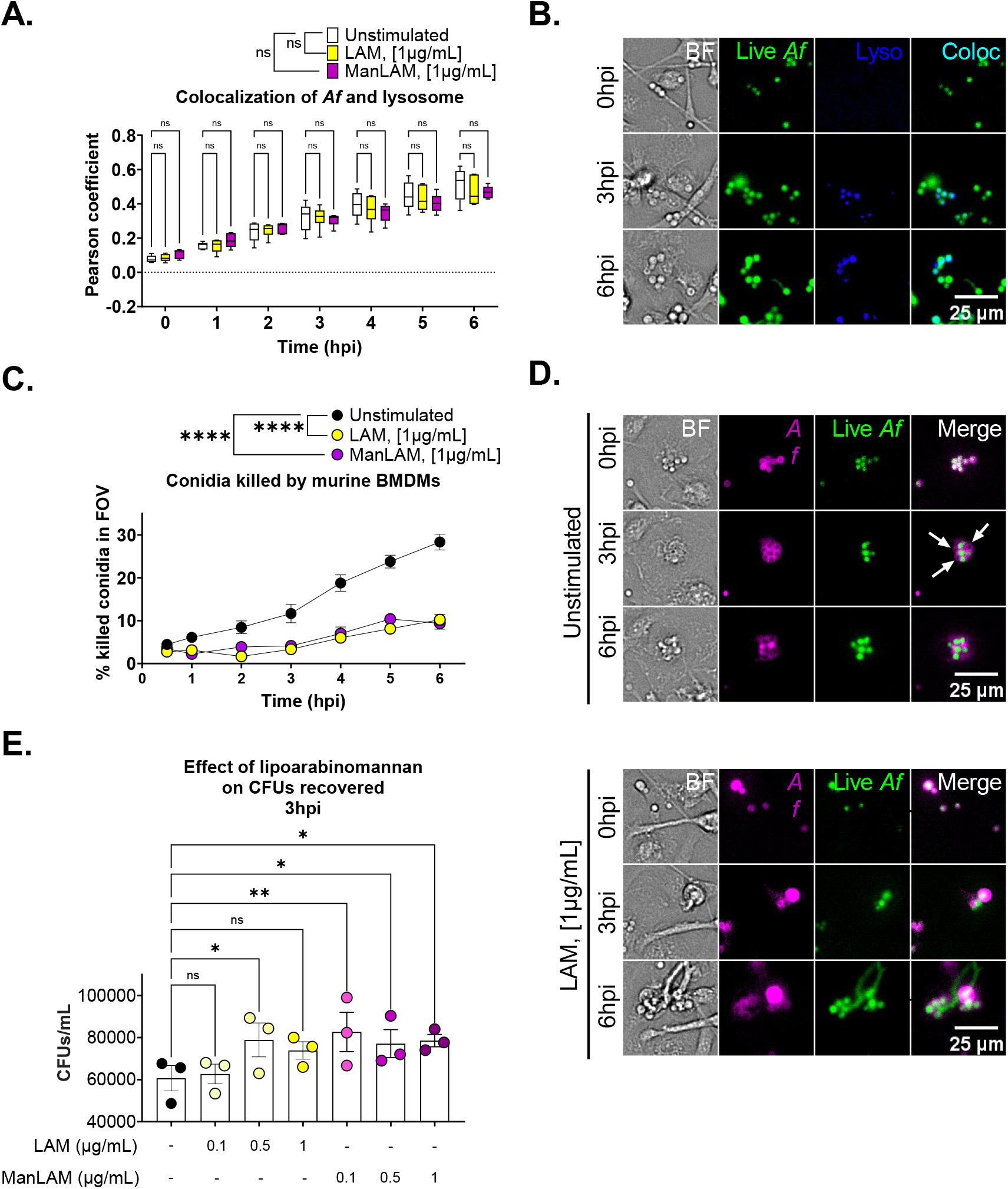
Mycobacterial ligands reduce macrophages killing rate of conidia. **A**. Pearson coefficient for *Af*-lysosome colocalization over time show increasing trend over time for all conditions without statistically significant differences between them. **B**. Image of colocalization analysis at 0, 3, and 6 hpi. **C**. Percentage of killed conidia (counted as single Alexa Fluor 633 positive signal) over time for US, LAM, and ManLAM. **D**. Comparative images of BMDMs killing *Af* conidia. White arrows indicate dead Af conidia (single Alexa Fluor 633 positive). **E**. CFUs results from lysed cells stimulate with LAM and ManLAM at 3hpi. Two-way ANOVA test for colocalization experiment and conidia killing assay. Box-plot centre line, median; +, mean, upper and lower quartiles; whiskers, 10^th^ and 90^th^ percentile. One-way ANOVA for CFUs. Statistical significance was tested on biological replicates (n=3). p value * < 0.05, ** < 0.01, *** < 0.001, **** < 0.0001

Nonetheless, using a FLuorescent Aspergillus REporter (FLARE) conidia (28), which allows the tracking in real-time of fungus clearance, a significant decrease in conidia killing was detected with both, LAM and ManLAM. By 2 hpi, LAM killed 1.7% vs 8.4% from unstimulated controls, which represents 80% less killing. At 6 hpi, LAM killed 10.2% of conidia vs 28.3% from unstimulated cells, or 74% less conidia. For ManLAM, the effect was similar. By 1 hpi, ManLAM cleared 2.5% vs 6.1% in controls. At 6 hpi, BMDMs stimulated with ManLAM cleared only 9.4% of conidia (Figure 1 C D). These findings were confirmed with CFUs counting. At 3 hpi, significant more CFUs were obtained from cells stimulated with LAM and ManLAM. LAM showed a dose effect, with statistically significant difference detected at 0.5 μg/mL but lost at 1 μg/mL. Similarly, ManLAM showed a stronger effect with 0.1 μg/mL than with 0.5 and 1 μg/mL (Figure 4 E).

## DISCUSSION

Here, we show that mycobacterial ligand, lipoarabinomannan, negatively modulates macrophage responses to *Af* conidia at early stages of infection. LAM increased cell death and IL-1β release, while LAM and ManLAM decreased the killing of *Af* conidia at early stages of infection in an equivalent manner. These results suggest that mycobacterial virulence factors can interfere effectively against early immune response towards *Af* spores.

*Af* adaptation to its environment and its metabolic versatility has been highlighted as a factor contributing to its success as an opportunistic pathogen (29). Furthermore, interference with metabolic pathways has been shown to modify its virulence significantly. For example, high availability of glucose and acetate, carbon energy sources, has been shown to change cell-wall components that enhance the resistance to oxidative stress (30). It is unknown if mycobacteria have the capacity to modulate fungal virulence. However, fungi have been shown to produce arabinases, which metabolizes sugars towards L-arabinose and the activation of the pentose-phosphate pathway (31). Whether if lipoarabinomannan can biochemically modulate *Af* fitness or development via induction of specific metabolic pathways is uncertain. Nonetheless, we showed that neither LAM or ManLAM produced any changes in the growth rate and germination of *Af* conidia, which suggests that our findings are most likely consequence of the modulation of macrophage responses and not due to early adaptations on fungi fitness, interpreted as growth rate and conidiation capacity (9).

BMDMs stimulation with LAM produced a significant increase in cell death. Mycobacteria have been shown to alter cell death pathways as a survival strategy (32,33). We suggested that these survival strategies prime macrophages to respond abnormally to *Af*. When macrophages were stimulated with LAM, more cells died faster, releasing significantly more IL-1β. *M. smegmatis*’ LAM is a strong TLR-2 agonist that leads to activation of NF-κβ. This has been shown to induce a hyperinflammatory response, characterised by increased expression of inflammasome components, and pyroptotic cell death (34,35). Findings with LAM were also consistent in J774A.1 cells (see Extended data), which showed increased cell death in a concentration-dependent manner and increased IL-1β release. This is consistent with NLRP3 inflammasome activation model (36), using LAM as signal 1 (priming), and *Af* as signal 2 (activation). Further investigation focused on the delivery of lysosome to the phagosome and conidia killing, since lipoarabinomannan has been shown to have the capacity to interfere with phagosome maturation and phago-lysosome fusion (37,38). Both, LAM and ManLAM showed significant decrease in the capacity to kill conidia within the first 6 hours of infection. However, this finding did not correlate with a reduced colocalization of lysosomes with *Af*. A 3hpi time-point was employed for counting CFUs from lysed macrophages, which corroborated the results using time-lapse microscopy.

Although, ManLAM binds preferentially to mannose receptor in macrophages (39), it also binds and activates TLR-2, eliciting inflammatory cytokine release (40,41). Interestingly, the stimulation of macrophages with 1 μg/mL of ManLAM led to reduced killing rate of conidia, without increased cell death or a differentiated inflammatory cytokine release. Higher concentrations of ManLAM, 10-20 µg/mL (42), have shown increased cytokine release, arrest of calcium-dependent processes, and apoptosis (43). However, we avoided these concentrations since they would have masqueraded priming.

Future studies should aim to characterise more deeply the broader effects of LAM on fungal-driven inflammation kinetically, and the impact on cell exhaustion. The failure to kill *Af* conidia at early stages of infection suggests that these mechanisms might be compromised. To corroborate the clinical significance of these findings, further testing using human monocyte-derived macrophages from patients with *Mtb* and atypical mycobacterial infections would be beneficial.

Taking into consideration that deficiency of NOD2 expression is protective for invasive *Af* infection (18), we could suggest that conditions that boost NOD2 activation lead to undesirable outcomes. NOD2 increases NF-κβ, which primes NLRP3 for activation (49–51). TLR-2 and NOD2 have been characterised as non-redundant systems in the recognition of *Mtb*, although their synergic response is necessary for the induction of proinflammatory cytokines (46).

We propose that mycobacteria’s virulence factors could shift the early immune response to conidia, since the inflammasome activation has been reported exclusively for hyphae (19,21). Moreover, in a keratitis model it has been reported that TLR-2 – NOD2 synergism provides an enhanced inflammatory cytokine response to conidia (47). By activating the TLR-2 signalling through the MDP anchor in lipoarabinomannan and MDP in mycobacterial cell wall NOD2 activation could be triggered. Theoretically, when *Af* encounters the macrophage, it can provide an additional stimulus for TLR-2. Via phagocytosis, *Af* conidia is recognized by NOD2, inducing further activation stimuli. Both, TLR-2 and NOD2 pathways meet downstream with activation of NF-κβ, leading to NLRP3 assembly, formation of the inflammasome, caspase-1 activation and pyroptotic cell death, with subsequent release of IL-1β and *Af* conidia to the extracellular space. Nonetheless, this will require further investigation.

In conclusion, our results indicates that lipoarabinomannan can modulate macrophage responses to *Af*, interfering with clearance of fungi. LAM, lipoarabinomannan without mannose cap, induces a hyperinflammatory response characterised by increased release of IL-1β and increased rate of macrophages death.

## AUTHOR CONTRIBUTION

Luis Edgardo Gonzales-Huerta

Roles: Conceptualization, Data Curation, Formal Analysis, Funding Acquisition, Investigation, Methodology, Project Administration, Software, Resources, Validation, Visualization, Writing – Original Draft Preparation, Writing – Review & Editing

Thomas Williams

Roles: Conceptualization, Methodology, Project Administration, Visualization, Writing – Review & Editing

Renad Aljohani

Roles: Conceptualization, Methodology, Project Administration, Visualization, Writing – Review & Editing

Brian Robertson

Roles: Conceptualization, Data Curation, Formal Analysis, Investigation, Methodology, Project Administration, Software, Resources, Validation, Visualization, Writing – Original Draft Preparation, Writing – Review & Editing

Carlton AW Evans

Roles: Conceptualization, Data Curation, Formal Analysis, Funding Acquisition, Investigation, Methodology, Project Administration, Software, Resources, Validation, Visualization, Writing – Original Draft Preparation, Writing – Review & Editing

Darius Armstrong-James

Roles: Conceptualization, Data Curation, Formal Analysis, Funding Acquisition, Investigation, Methodology, Project Administration, Software, Resources, Supervision, Validation, Visualization, Writing – Original Draft Preparation, Writing – Review & Editing

## EXTENDED DATA

**Extended Data 1.**
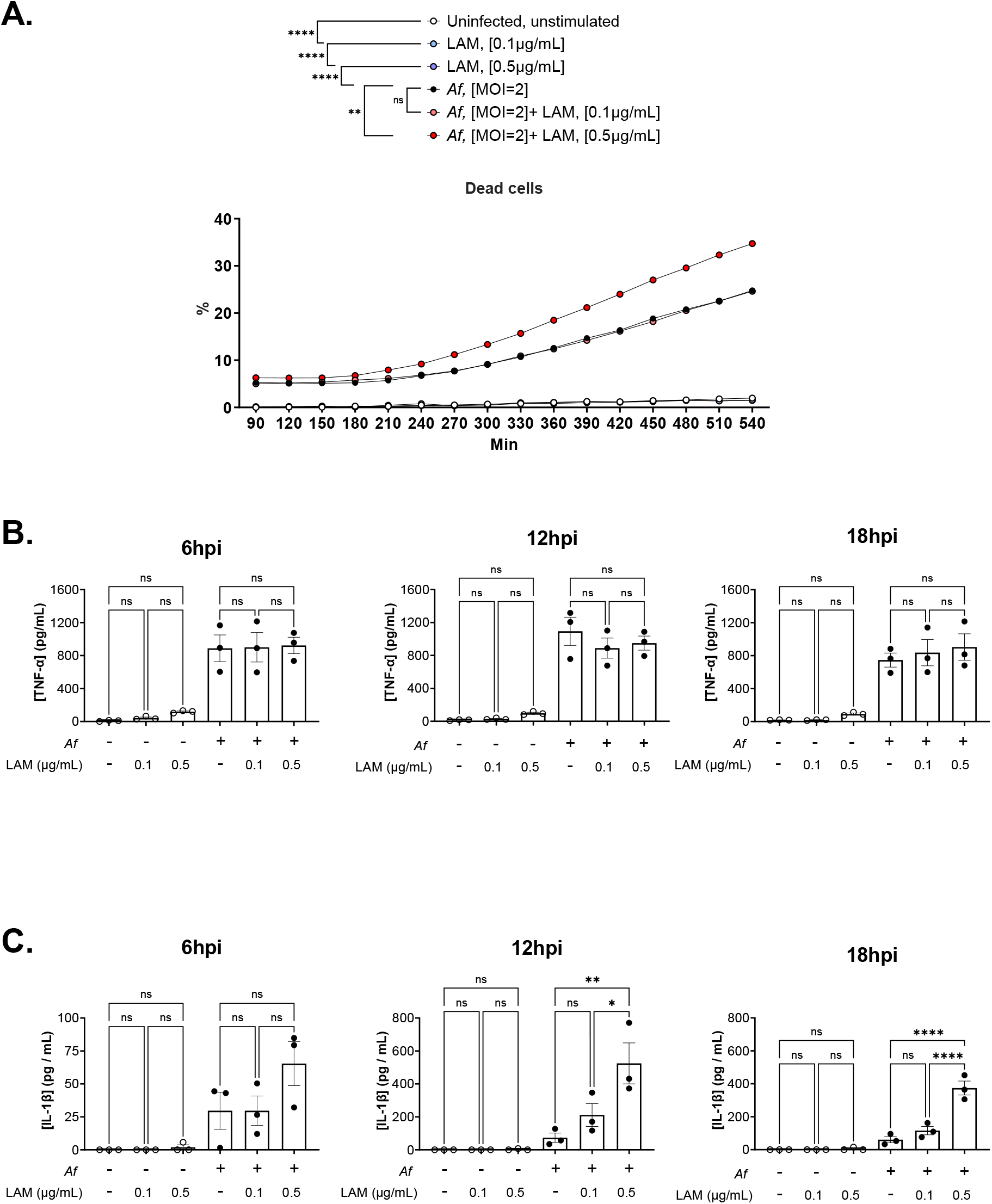
LAM increases cell death and IL-1β release in J774A.1 cells infected with *Af*. **A**. Time-lapse fluorometric assay using PI as a cell death marker. Percentages were calculated as relative to control with 0.1% TX100. **B**. Concentration of TNF-α in supernatant from J774A.1 cells infected with *Af* for 6, 12, and 18 hpi. **C**. Concentration of IL-1β in supernatant from cells infected with *Af* for 6, 12, and 18 hpi. Both cytokines were measured simultaneously from the same sample. One-way ANOVA was performed on 3 experimental replicates (n=3). Error bars in SEM.

**Extended Data 2.**
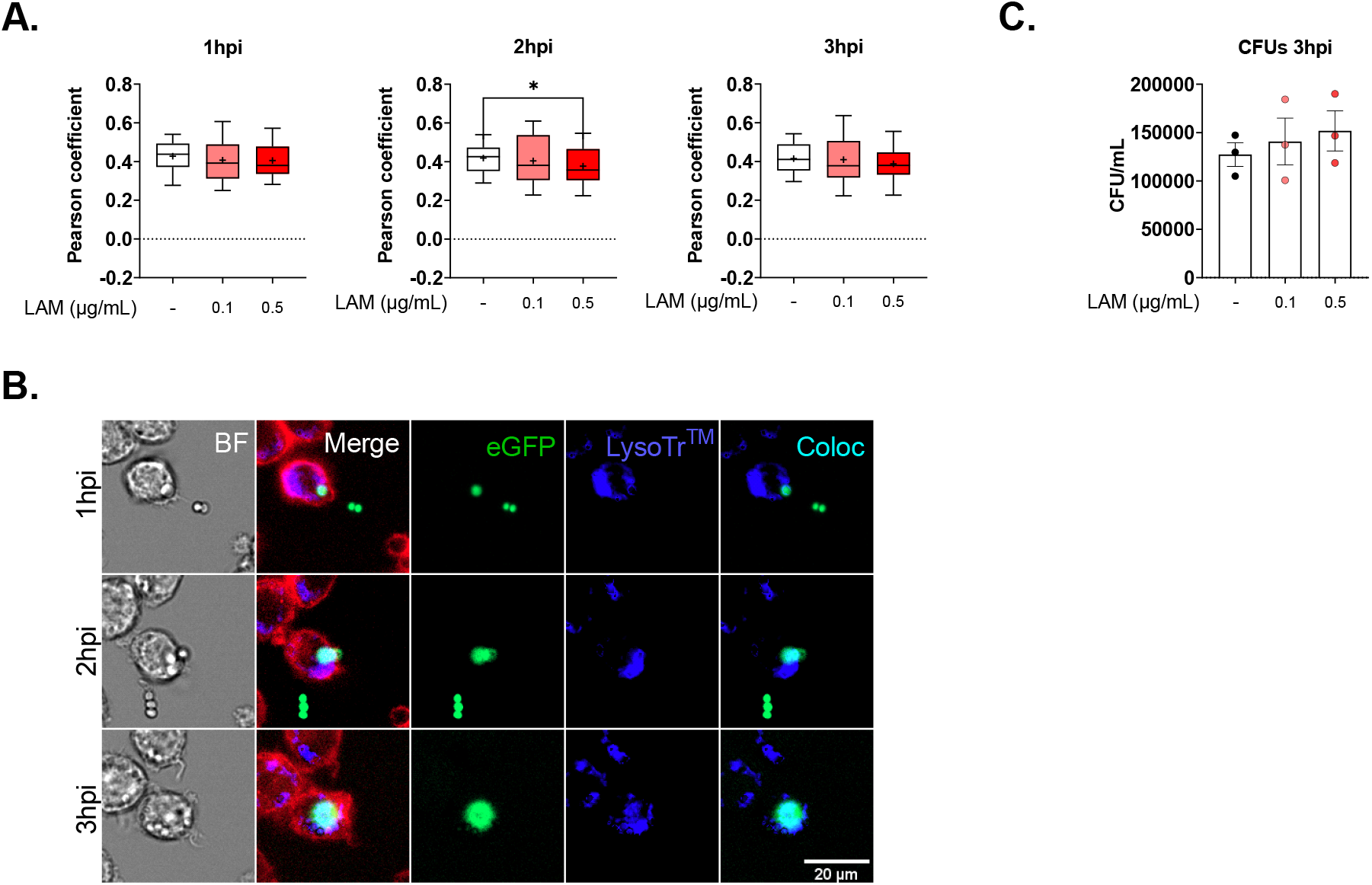
LAM reduced phagolysosome fusion at early stages of infection in J774A.1 cells. **A**. Colocalization of eGFP-*Af* and Lysotracker™ Blue DND-22 at 1, 2, and 3 hpi presented as Pearson’ Coefficient. Samples pooled from 3 experimental replicates for n=81 cells per condition. **B**. Representative images widefield time-lapse fluorescence microscopy showing J774A.1 cells, stained with CD45-AF700 (red) stimulated with 0.5µg/mL of LAM infected with eGFP-*Af* (green) and stained for lysosome LysoTracker™ Blue DND-22 (blue). Colocalized areas can be identified in cyan. **C**. CFUs obtained from J774A.1 cells lysed at 3 hpi. Two-way ANOVA test for colocalization experiment and one-way ANOVA for CFUs. Box-plot centre line, median; +, mean, upper and lower quartiles; whiskers, 10^th^ and 90^th^ percentile.

